# Cross-neutralization of Omicron BA.1 against BA.2 and BA.3 SARS-CoV-2

**DOI:** 10.1101/2022.03.30.486409

**Authors:** Jing Zou, Chaitanya Kurhade, Hongjie Xia, Mingru Liu, Xuping Xie, Ping Ren, Pei-Yong Shi

## Abstract

The Omicron SARS-CoV-2 has three distinct sublineages, among which sublineage BA.1 is responsible for the initial Omicron surge and is now being replaced by BA.2 world-wide, whereas BA.3 is currently at a low frequency. The ongoing BA.1-to-BA.2 replacement underscores the importance to understand the cross-neutralization among the three Omicron sublineages. Here we tested the neutralization of BA.1-infected human sera against BA.2, BA.3, and USA/WA1-2020 (a strain isolated in late January 2020). The BA.1-infected sera neutralized BA.1, BA.2, BA.3, and USA/WA1-2020 SARS-CoV-2s with geometric mean titers (GMTs) of 445, 107, 102, and 16, respectively. Thus, the neutralizing GMTs against heterologous BA.2, BA.3, and USA/WA1-2020 were 4.2-, 4.4-, and 28.4-fold lower than the GMT against homologous BA.1, respectively. These findings have implications in COVID-19 vaccine strategy.

## Main text

Since the emergence of severe acute respiratory syndrome coronavirus 2 (SARS-CoV-2) in late 2019, the virus has evolved to increase viral transmission and immune evasion. The World Health Organization (WHO) has so far designated 5 variants of concern (VOC), including Alpha, Beta, Gamma, Delta, and Omicron. The most recently emerged Omicron variant has 3 distinct sublineages: BA.1, BA.2, and BA.3. BA.1 was first identified in South Africa in November 2021. BA.1 and its derivative BA.1.1 (containing an extra R346K substitution in the spike of BA.1) caused the initial surges of Omicron around the world. Subsequently, the frequency of BA.2 increased steeply, replacing BA.1 in many parts of the world. In the USA, the frequency of BA.2 increased from 0.4% to 54.9% between January 22 and March 24, 2022 (https://covid.cdc.gov/covid-data-tracker/#variant-proportions). Compared with BA.1, BA.2 did not seem to cause more severe disease^1^, but may increase viral transmissible by ∼30%^2^. As of March 30, 2022, the frequency BA.3 remained low in the GISAID database (https://www.gisaid.org/). All three sublineages of Omicron could significantly evade vaccine-elicited neutralization, among which BA.3 exhibited the greatest reduction^3,4^. In addition, Omicron BA.1 could efficiently evade non-Omicron SARS-CoV-2 infection-elicited neutralizationzou^5^. The increased transmissibility and immune evasion of the Omicron variant may be responsible for the replacement of VOC from the previous Delta to the current Omicron. Since many unvaccinated individuals were infected by BA.1 during the initial Omicron surge, it is important to examine the cross-neutralization of BA.1 infection against BA.2, BA.3, and other variants. Such laboratory information is essential to guide vaccine strategy and public health policy.

To examine the cross-neutralization among the three Omicron sublineages, we collected 20 human sera from unvaccinated patients who were infected with Omicron BA.1 (**Table 1**). The genotype of infecting virus was verified for each patient by Sanger sequencing. The sera were collected on day 8 to 62 after positive RT-PCR test. The serum panel was measured for neutralization against four recombinant SARS-CoV-2s (**Figure 1A**): USA/WA1-2020 (wild-type) and three chimeric USA/WA1-2020 bearing the full-length spike protein from Omicron BA.1 (GISAID EPI_ISL_6640916), BA.2 (GISAID EPI_ISL_6795834.2), or BA.3 (GISAID EPI_ISL_7605591). The spike proteins of the three Omicron sublineages have distinct amino acid mutations, deletions, and insertions (**Figure 1A**). To facilitate neutralization testing, an mNeonGreen (mNG) reporter was engineered into the four viruses, resulting in wild-type, BA.1-, BA.2-, and BA.3-spike mNG SARS-CoV-2s. The construction and characterization of the four mNG SARS-CoV-2s were recently reported^4^. Using an mNG-based fluorescent focus-reduction neutralization test (FFRNT), we determined the neutralizing geometric mean titers (GMTs) of the sera against wild-type, BA.1-, BA.2-, and BA.3-spike mNG SARS-CoV-2s to be 445, 107, 102, and 16, respectively (**Fig. 1B**). Thus, the neutralizing GMTs against heterologous BA.2-spike, BA.3-spike, and wild-type viruses were 4.2-, 4.4-, and 28.4-fold lower than the GMT against the homologous BA.1-spike virus, respectively (**Fig. 1B**). Consistently, all sera neutralized BA.1-spike virus at neutralizing titers of ≥80, whereas 12 out of 20 sera did not neutralize the wild-type USA/WA1-2020 (defined as 10 for plot and calculation purposes; **Fig. 1B** and **Table 1**). Notably, 2 sera neutralized BA.2-spike virus more efficiently than the BA.1-spike virus (indicated by symbol * in **Fig. 1C**). Collectively, the results support two conclusions. First, BA.1 infection elicited similar levels of cross-neutralization against BA.2 and BA.3, although at a decreased efficiency that was 4.2-to 4.4-fold lower than that against BA.1. This result is in contrast with the neutralization results from vaccinated sera (collected at 1 month after three doses of Pfizer/BioNTech’s BNT162b2 vaccine) which neutralized BA.1 and BA.2 much more efficiently than BA.3^4^. Second, the neutralization of BA.1-infected sera against USA/WA1-2020 were 6.7- and 6.4-fold lower than that against Omicron BA.2 and BA.3, respectively. The results indicate the antigenic distinctions among different variant spikes, which must be carefully considered when deciding to switch the vaccine sequence to new variants^6^. If future variants are Omicron decedents, switch vaccine sequence to an Omicron spike is conceptually attractive.

**Table 1.** Serum information and FFRNT_50_ values.

| Serum ID | Age (years) | Gender (F/M) | Ethnicity | *FFRNT<sub>50</sub> |  |  |  | Serum collection time (days post positive RT-PCR test) |
| --- | --- | --- | --- | --- | --- | --- | --- | --- |
|  |  |  |  | USA-WA1/2020 | BA.1-spik<br>e<br>virus | BA.2-s<br>pike<br>virus | BA.3-s<br>pike<br>virus |  |
| 1 | 29 | M | Hispanic | ^10 | 80 | 160 | 20 | 26 |
| 2 | 41 | F | White | 10 | 80 | 113 | 40 | 33 |
| 3 | 84 | M | White | 14 | 113 | 40 | 14 | 21 |
| 4 | 14 | M | Black | 10 | 113 | 28 | 28 | 16 |
| 5 | 35 | F | Black | 10 | 160 | 40 | 40 | 16 |
| 6 | 28 | F | Black | 10 | 160 | 20 | 20 | 28 |
| 7 | 9 | F | Hispanic | 10 | 160 | 57 | 57 | 43 |
| 8 | 64 | M | Black | 10 | 160 | 80 | 20 | 40 |
| 9 | 84 | M | White | 20 | 226 | 160 | 226 | 40 |
| 10 | 5 | F | Hispanic | 10 | 320 | 28 | 40 | 26 |
| 11 | 1 | F | Hispanic | 10 | 320 | 113 | 160 | 56 |
| 12 | 57 | F | White | 10 | 453 | 28 | 80 | 35 |
| 13 | 75 | M | Hispanic | 160 | 453 | 10 | 20 | 17 |
| 14 | 26 | F | Hispanic | 20 | 640 | 453 | 160 | 62 |
| 15 | 55 | M | White | 10 | 905 | 40 | 113 | 29 |
| 16 | 74 | F | <sup>\$</sup>NA | 10 | 1280 | 226 | 320 | 8 |
| 17 | 71 | F | Hispanic | 28 | 2560 | 453 | 640 | 32 |
| 18 | 63 | M | Hispanic | 113 | 2560 | 1280 | 1810 | 15 |
| 19 | 78 | F | Hispanic | 10 | 5120 | 640 | 1280 | 16 |
| 20 | 84 | M | White | 40 | 14482 | 2560 | 2560 | 13 |
| <sup>#</sup>GMT | 34 | - | - | 16 | 445 | 107 | 102 | 25 |
| <sup>†</sup>95% CI | 20-58 | - | - | 11-24 | 225-881 | 54-215 | 48-218 | 20-32 |

\*Individual FFRNT<sub>50</sub> value is the geometric mean of duplicate plaque assay results.
^FFRNT<sub>50</sub> of <20 was treated as 10 for plot purpose and statistical analysis.
<sup>\$</sup>NA: not available.
<sup>#</sup>Geometric mean neutralizing titers (GMT).
<sup>†</sup>95% confidence interval (95% CI) for the GMT.

**Figure 1.**
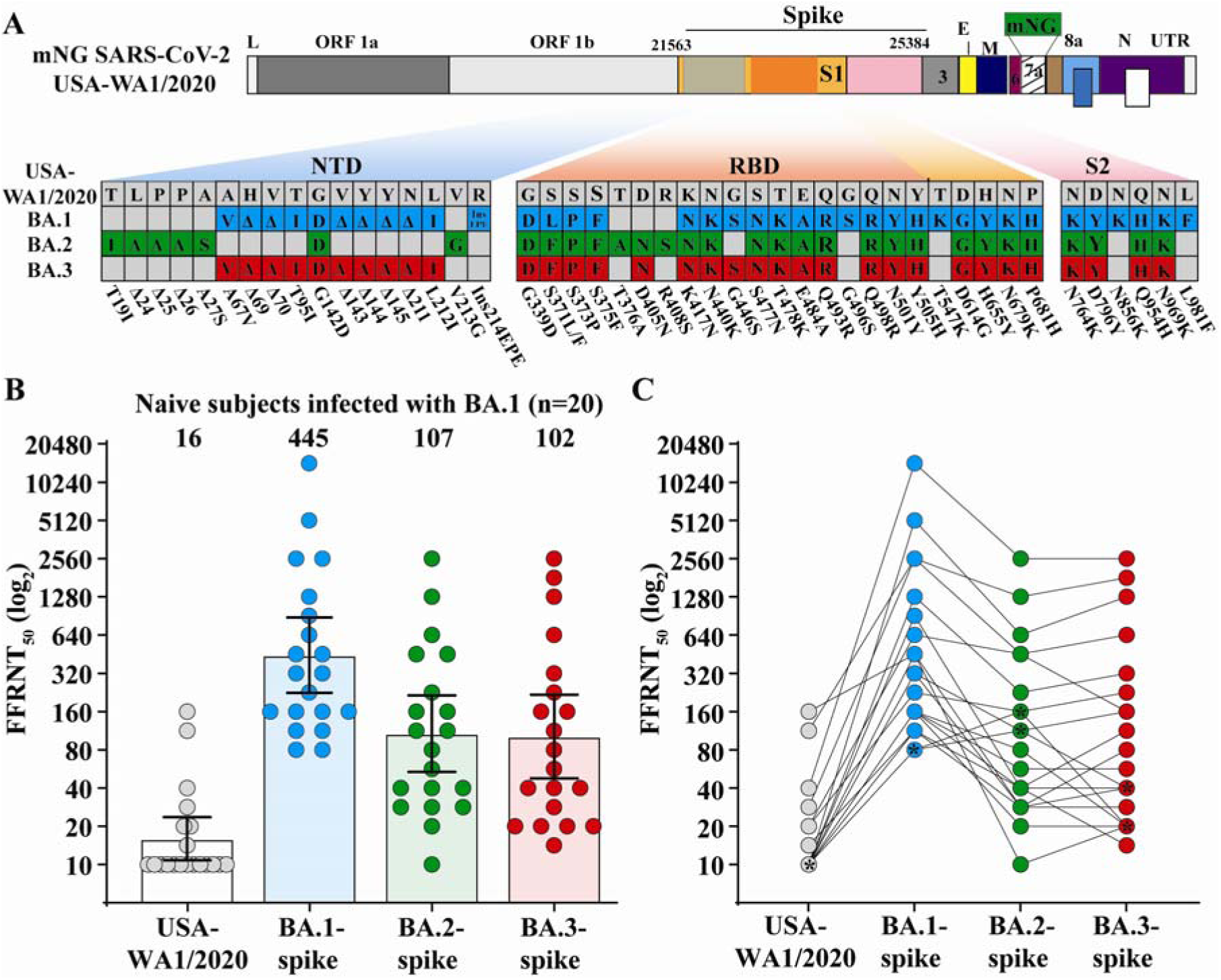
Cross-neutralization of human sera from unvaccinated individuals who were infected with Omicron BA.1 SARS-CoV-2. (A) Omicron BA.1-, BA.2-, and BA.3-spike mNG SARS-CoV-2s. The full-length spike gene from Omicron BA.1, BA.2, or BA.3 was engineered into an mNG USA-WA1/2020 SARS-CoV-2. The mNG gene was engineered at the open-reading-frame-7 of the viral genome. Amino acid mutations, deletions, and insertions (Ins) are indicated for BA.1, BA.2, and BA.3 spikes in reference to the USA-WA1/2020 spike. L: leader sequence; ORF: open reading frame; NTD: N-terminal domain of S1; RBD: receptor binding domain of S1; S: spike glycoprotein; S1: N-terminal furin cleavage fragment of S; S2: C-terminal furin cleavage fragment of S; E: envelope protein; M: membrane protein; N: nucleoprotein; UTR: untranslated region. (B) Scatterplot of neutralization titers. A panel of 20 human sera collected from Omicron BA.1-infected individuals were tested for the 50% fluorescent focus-reduction neutralization titers (FFRNT_50_) against recombinant USA-WA1/2020, Omicron BA.1-, BA.2-, and BA.3-spike mNG SARS-CoV-2s. The neutralization titer for each virus was determined in duplicates. The serum information and FFRNT_50_ values are summarized in **Table 1**. Each data point represents the geometric mean FFRNT_50_ obtained with a serum specimen against the indicated virus. The bar heights and the numbers above indicate geometric mean titers (GMTs). Error bars indicate the 95% confidence interval (CI) of the GMTs. Statistical analysis was performed using the Wilcoxon matched-pairs signed-rank test. *P* values of the GMT against BA.1-spike and USA-WA1/2020, BA.2-spike, or BA.3 spikes are all <0.0001. (C) FFRNT_50_ values with connected lines for individual sera. Two sera exhibiting slightly higher FFRNT_50_s against BA.2 virus than that against BA.1-spike SARS-CoV-2 are indicated by symbol * (serum ID 1 and 2 in **Table 1**).

Emerging evidence supports a vaccine booster strategy to minimize the health risk of the ongoing Omicron surge. First, 2 doses of BNT162b2 vaccine are inefficient to elicit robust neutralization against Omicron variant, whereas 3 doses of BNT162b2 produces robust neutralization against Omicron. Although Omicron-neutralizing activity remains robust for up to 4 months^3^, the durability of such neutralization beyond 4 months after dose 3 remains to be determined. The latter result, together with the real-world vaccine effectiveness, are required to guide the timing of dose 4 vaccine. Second, non-Omicron SARS-CoV-2 infection does not elicit robust neutralization against Omicron variant^5^, suggesting that previously infected individuals should be vaccinated to mitigate the health threat from Omicron. The cross-neutralization of BA.1-infected sera against BA.2 and BA.3 suggests the recent BA.1-infected individuals are likely to be protected against the ongoing BA.2 surge. Third, vaccine-mediated T cell immunity and non-neutralizing antibodies that mediate antibody-dependent cytotoxicity could also confer protection against severe COVID-19. After vaccination or infection, the majority of T cell epitopes are highly preserved against Omicron spikes^7^. In agreement with this notion, 3 doses of BNT162b2 conferred efficacy against Omicron disease, but the protection wanes over time, with overall efficacy remaining high up to 6 months after dose 3^8-12^. The real-world vaccine effectiveness and laboratory studies will guide vaccine booster strategy to achieve optimal breadth and duration of protection.

## Methods

### Recombinant Omicron spike mNG SARS-CoV-2s

The construction and characterization of recombinant Omicron BA.1-, BA.2-, and BA.3-spike mNG SARS-CoV-2s were recently reported^4^. The BA.1, BA.2, and BA.3 spike sequences were derived from GISAID EPI_ISL_6640916, EPI_ISL_6795834.2, and EPI_ISL_7605591, respectively. Passage 1 (P1) virus stocks were produced from infectious cDNA clones of corresponding viruses^13,14^. The P1 viruses were used for neutralization testing throughout the study. The spike gene from each P1 virus was sequenced to ensure no undesired mutations. Equivalent specific infectivities, defined by the genomic RNA-to-FFU (fluorescent focus-forming unit) ratios, were confirmed for individual recombinant P1 virus stocks, as previously reported^4^.

### Serum specimens

The research protocol regarding the use of human serum specimens was reviewed and approved by the University of Texas Medical Branch (UTMB) Institutional Review Board (IRB number 20-0070). The de-identified human sera from unvaccinated patients who were infected by Omicron sublineage BA.1 were heat-inactivated at 56°C for 30 min before neutralization testing. The genotype of infecting virus was verified by the molecular tests with FDA’s Emergency Use Authorization and Sanger sequencing. The serum information is presented in **Table 1**.

### Fluorescent focus reduction neutralization test

Neutralization titers of sera were measured by fluorescent focus reduction neutralization test (FFRNT) using the USA-WA1/2020, BA.1-, BA.2-, and BA.3-spike mNG SARS-CoV-2s. The FFRNT protocol was reported previously^5^. Briefly, Vero E6 cells were seeded to 96-well plates at 2.5×10^4^ per well (Greiner Bio-one™). On the following day, heat-inactivated sera were 2-fold serially diluted in culture medium with the first dilution of 1:20 (final dilutions ranging from 1:20 to 1:20,480). The diluted serum was incubated with 100-150 FFUs of indicated mNG SARS-CoV-2s at 37°C for 1 h. Afterwards, the serum-virus mixtures were loaded onto the pre-seeded Vero E6 cell monolayer in 96-well plates. After 1 h infection, the inoculum was aspirated and overlay medium (100 μl supplemented with 0.8% methylcellulose) was added to each well. After incubating the plates at 37°C for 16-18 h, raw images of mNG foci were acquired using Cytation™ 7 (BioTek). The foci in each well were counted and normalized to the no-serum-treated controls to calculate infection rates. The FFRNT_50_ value was defined as the minimal serum dilution that suppressed >50% of fluorescent foci. The neutralization titer of each serum was determined in duplicates, and the geometric mean was presented. FFRNT_50_ of <20 was treated as 10 for plot purpose and statistical analysis. **Tables 1** summarizes the FFRNT_50_ results.

### Statistics

The nonparametric Wilcoxon matched-pairs signed rank test was used to analyze the statistical significance in **Figure 1B**.

### Data availability

The data that support the findings of this study are available from the corresponding authors upon request.

## Acknowledgments

We thank colleagues at the University of Texas Medical Branch (UTMB) for helpful discussions. We thank Michael L O’Rourke from the Information System Department at UTMB for assisting with electronic medical record systems. P.-Y.S. was supported by NIH grants HHSN272201600013C, AI134907, AI145617, and UL1TR001439, and awards from the Sealy & Smith Foundation, the Kleberg Foundation, the John S. Dunn Foundation, the Amon G. Carter Foundation, the Gilson Longenbaugh Foundation, and the Summerfield Robert Foundation.

## Author contributions

Conceptualization, X.X., P.R., P.-Y.S.; Methodology, J.Z., C.K., M.L., X.X.,

P.R., P.-Y.S; Investigation, J.Z., C.K., M.L., X.X., P.R., P.-Y.S; Resources, C.K.,

M.L., X.X., P.R., P.-Y.S; Data Curation, J.Z., C.K., X.X., P.R., P.-Y.S.; Writing-Original Draft, J.Z., H.X., X.X., R.R., P.-Y.S; Writing-Review & Editing, X.X.,

P.R., P.-Y.S.; Supervision, X.X., P.R., P.-Y.S.; Funding Acquisition, P.-Y.S.

## REFERENCES

1. Nicole Wolter, Waasila Jassat, DATCOV-Gen author group, Anne von Gottberg, Cohen C. Clinical severity of Omicron sub-lineage BA.2 compared to BA.1 in South Africa. MedRxiv 2022: https://doi.org/10.1101/2022.02.17.22271030.

2. Frederik Plesner Lyngse, Carsten Thure Kirkeby, Matthew Denwood, et al. Transmission of SARS-CoV-2 Omicron VOC subvariants BA.1 and BA.2: Evidence from Danish Households. BioRxiv 2022:doi: https://doi.org/10.1101/2022.01.28.22270044.

3. Xia H, Zou J, Kurhade C, et al. Neutralization and durability of 2 or 3 doses of the BNT162b2 vaccine against Omicron SARS-CoV-2. Cell Host Microbe 2022.

4. Kurhade C, Zou J, Xia H, et al. Neutralization of Omicron BA.1, BA.2, and BA.3 SARS-CoV-2 by 3 doses of BNT162b2 vaccine. BioRxiv 2022:https://biorxiv.org/cgi/content/short/2022.03.24.485633v1.

5. Zou J, Xia H, Xie X, et al. Neutralization against Omicron SARS-CoV-2 from previous non-Omicron infection. Nat Commun 2022;13:852.

6. Liu Y, Liu J, Zou J, et al. Distinct neutralizing kinetics and magnitudes elicited by different SARS-CoV-2 variant spikes. bioRxiv 2021.

7. Redd AD, Nardin A, Kared H, et al. Minimal Crossover between Mutations Associated with Omicron Variant of SARS-CoV-2 and CD8(+) T-Cell Epitopes Identified in COVID-19 Convalescent Individuals. mBio 2022:e0361721.

8. Ferdinands JM, Rao S, Dixon BE, et al. Waning 2-Dose and 3-Dose Effectiveness of mRNA Vaccines Against COVID-19-Associated Emergency Department and Urgent Care Encounters and Hospitalizations Among Adults During Periods of Delta and Omicron Variant Predominance -VISION Network, 10 States, August 2021-January 2022. MMWR Morb Mortal Wkly Rep 2022;71:255–63.

9. Chemaitelly H, Abu-Raddad LJ. Waning effectiveness of COVID-19 vaccines. Lancet 2022;399:771–3.

10. Tartof SY, Slezak JM, Puzniak L, et al. BNT162b2 (Pfizer–Biontech) mRNA COVID-19 Vaccine Against Omicron-Related Hospital and Emergency Department Admission in a Large US Health System: A Test-Negative Design. SSRN 2022:https://ssrn.com/abstract=4011905 or http://dx.doi.org/10.2139/ssrn.

11. Andrews N, Stowe J, Kirsebom F, et al. Covid-19 Vaccine Effectiveness against the Omicron (B.1.1.529) Variant. N Engl J Med 2022.

12. Agency UHS. COVID-19 vaccine surveillance report – Week 9, 3 March 2022. 2022:https://assets.publishing.service.gov.uk/government/uploads/system/uploads/attachment_data/file/1058464/Vaccine-surveillance-report-week-9.pdf.

13. Xie X, Muruato A, Lokugamage KG, et al. An Infectious cDNA Clone of SARS-CoV-2. Cell Host Microbe 2020;27:841–8 e3.

14. Xie X, Lokugamage KG, Zhang XV M.N., Muruato AE, Menachery VD, Shi P-Y. Engineering SARS-CoV-2 using a reverse genetic system. Nature Protocols 2021;16:1761–84.

